# Neural graft elimination via dual safety switch compromises therapeutic recovery after stroke

**DOI:** 10.64898/2026.05.29.728814

**Authors:** Beatriz Achón Buil, Nora H. Rentsch, Rebecca Z. Weber, Carmen Helfenstein, Chantal Bodemann, Kathrin J Zürcher, Siri Peter, Melanie Generali, Christian Tackenberg, Ruslan Rust

**Author notes:** **Correspondence:** Ruslan Rust, Ph.D. Assistant Professor, The Zilkha Neurogenetic Institute, Department of Physiology and Neuroscience, Keck School of Medicine of the University of Southern California 1501 San Pablo Street, Los Angeles, CA 90033; Christian Tackenberg, Ph.D. Scientific Head of Division Neurodegeneration Institute for Regenerative Medicine • IREM University of Zurich, Wagistrasse 12, 8952 Schlieren, Switzerland.

## Abstract

Safety switch systems are increasingly incorporated into stem cell therapies to enable selective graft elimination in the event of adverse outcomes. Yet how removing transplanted cells affects the underlying pathology remains largely unexplored. Here, we show that iPSC-derived neural stem cells (NSCs) co-transduced with herpes simplex virus thymidine kinase (HSV-TK) and inducible caspase-9 (iC9) are more efficiently ablated by combined ganciclovir (GCV) and chemical inducer of dimerization (CID) treatment than either single switch alone, under both proliferating and differentiating conditions. Using bioluminescence imaging to track graft survival longitudinally, we demonstrate that dual safety switch NSC (DS-NSC) grafts are selectively eliminated by GCV and CID administration in immunodeficient mice following photothrombotic stroke. The combination treatment initiated on day 8 post-transplantation resulted in reduced graft volume, lower NSC proliferation, and persistently reduced bioluminescence signal compared to mice receiving only solvent. Histological analysis revealed that combination treatment was associated with larger stroke lesions, higher IBA1 fluorescence intensity, and reduced vascular density in the ischemic border zone. Mice receiving GCV and CID showed persistently elevated contralateral hindlimb error rates on a ladder walk task from day 28 post-stroke onward compared to solvent-treated mice. These findings demonstrate that while DS-NSC elimination is feasible, the pharmacological activation of the safety switch system is associated with impaired functional stroke recovery.

## Introduction

Suicide gene therapy, originally developed to selectively eliminate cancer cells,^1^ has been adapted as a “safety switch” for stem cell therapies, enabling graft removal in the event of adverse outcomes such as graft-vs-host disease.^2^ For neurological disorders, such as stroke, stem cell therapy represents the main therapeutic avenue for functional recovery.^3^ However, the safety profile must be guaranteed before clinical translation,^4^ especially considering a potential future use of engineered cells, such as immune-evasive stem cells, which carry the risk of forming hypoimmunogenic tumors and becoming viral reservoirs.^5,6^

Among safety switch systems, enzyme/prodrug and inducible dimerization approaches are most suitable for intracerebral applications, as other strategies like antibody-mediated systems are hindered by the blood-brain barrier, which prevents antibodies from reaching and eradicating transplanted stem cells.^7,8^ The most extensively studied enzyme/prodrug system is herpes simplex virus thymidine kinase (HSV-TK), which converts ganciclovir (GCV) or its analogs into toxic metabolites that block DNA synthesis, thereby triggering necrotic and apoptotic processes.^9^ However, HSV-TK is immunogenic and exhibits a bystander effect, which refers to the ability to induce apoptosis in cells lacking the suicide gene by the diffusion of the converted toxic metabolite.^10,11^ Inducible caspase-9 (iC9) offers a complementary approach as it is non-immunogenic, its activator, chemical inducer of dimerization (CID), is bio-inert, and its activity is independent of proliferation.^12,13^

Yet, translating these systems clinically requires addressing their distinct limitations. In the case of HSV-TK, concerns regarding GCV toxicity have driven the development of safer prodrugs (e.g., brivudine, penciclovir) and more sensitive HSV-TK variants (e.g., SR39h, A168H point mutations).^14–16^ For iC9, the challenge lies in the heterogeneous killing efficiency, often caused by overexpression of BCL2, low initial levels of iC9, or low X-linked inhibitor of apoptosis protein (XIAP)/caspase-3 ratio.^17,18^ Strategies to overcome this issue include isolating an iC9 homogenous population sensitive to CID,^17^ or combining CID with XIAP inhibitors at a later treatment round.^18^

To further tune cell death induction, the combination of different safety switch systems has also been explored. For instance, the sequential activation of rapamycin-induced caspase-9 and HSV-TK safety switches in mesenchymal stem cells (MSCs) has been proven to be safe and to provide a failsafe mechanism that achieves higher killing efficiency.^19^ Additionally, co-transduction of Jurkat cells with rapamycin-sensitive iC9 and iC8 activatable by CID were more efficiently ablated with the combination treatment than single safety switches.^20^ Hence, a great effort has been put into improving the killing efficiency of safety switch systems; however, the effects that their ablation may cause remain largely unknown. Does the removal of the safety switch stem cells trigger a reaction that exacerbates the original pathology which was meant to be treated? Or are the initial benefits sufficient to justify proceeding with stem cell injection, even if the graft must be removed later on?

Here, we investigated the effects of ablating iPSC-derived neural stem cells (NSCs) in mice undergoing stroke induction*. In vitro,* NSCs co-transduced with the dual switch HSV-TK and iC9 (DS-NSCs) were more efficiently removed with the combination treatment of GCV and CID than single switches. *In vivo*, mice receiving DS-NSCs on day 7 post-stroke and treated a week later with GCV and CID presented smaller grafts and lower NSC proliferation rates than solvent-treated mice. Moreover, graft removal resulted in bigger stroke lesions, higher inflammation, lower vascular density, and less motor recovery from day 28 onwards. Collectively, our results show that the removal of NSC grafts from the stroke lesion although feasible may compromise functional recovery.

## Results

### Selective removal of NSCs expressing single or dual safety switch systems *in vitro*

To specifically ablate and longitudinally track grafted cells, we firstly generated two lentiviral plasmids containing either iC9 or HSV-TK together with red firefly luciferase (rFluc) under human elongation factor-1 alpha (EF1a) promoter (**Fig. 1A**). We included the hygromycin resistance gene to the iC9 construct and both GFP and puromycin resistance genes to the HSV-TK plasmid for subsequent selection. In addition, as a negative control, we used a lentiviral vector containing the same reporter and selectable genes as HSV-TK, but without any suicide gene (EF1a-rFluc-GFP_mPGK-Puro). NSCs were then transduced with iC9, HSV-TK, both (DS), or only rFluc lentiviral particles. NSCs containing the HSV-TK plasmid were sorted to isolate cells expressing high GFP levels and subsequently, all cell lines were selected with the corresponding antibiotic.

**Figure 1.**
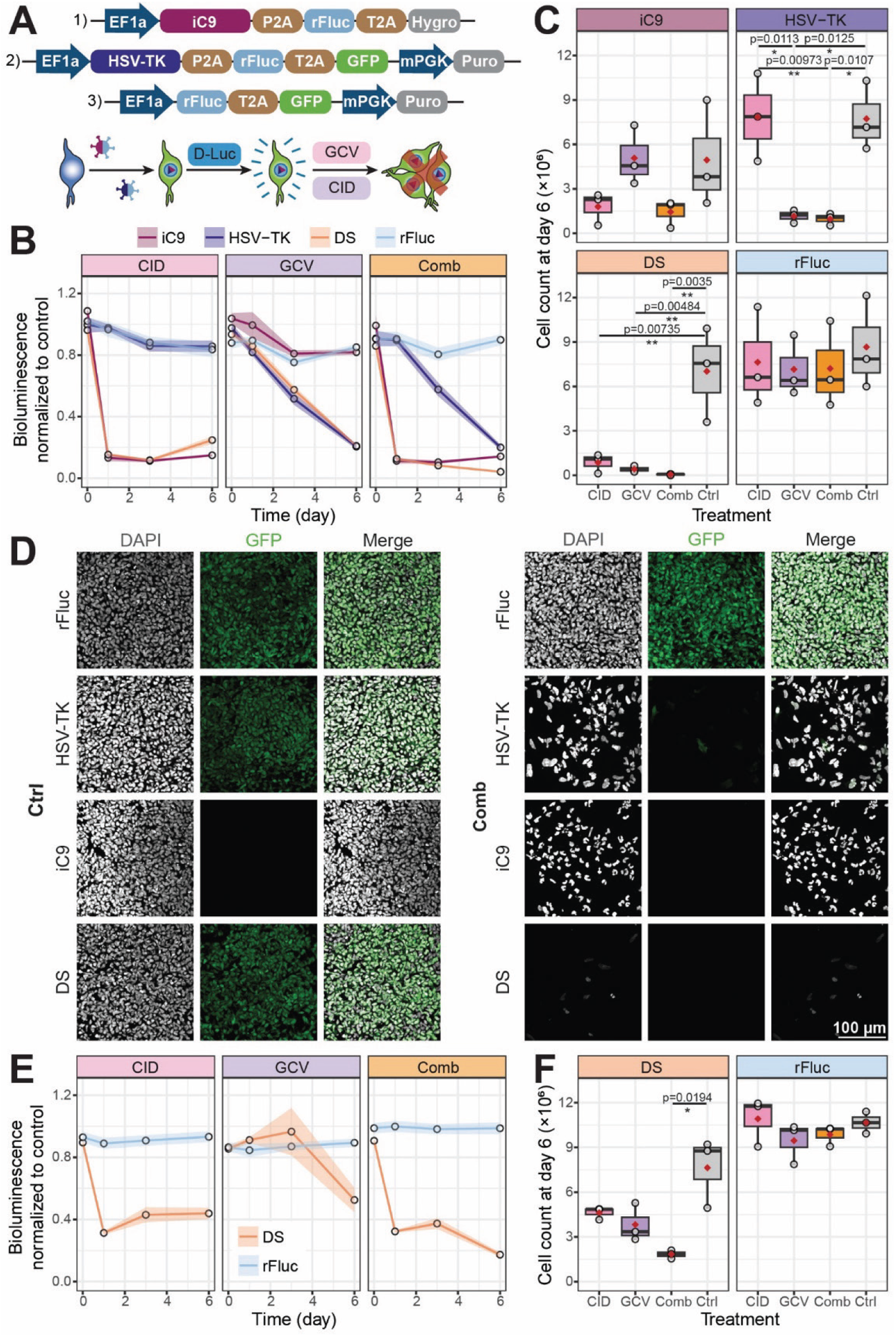
Removal of safety switch system neural stem cells (NSCs) *in vitro*. (**A**) We developed three lentiviral constructs containing 1) inducible caspase-9 (iC9), 2) herpes simplex virus thymidine kinase (HSV-TK), and/or 3) red firefly luciferase (rFluc) under human elongation factor-1 alpha (EF1a) promoter together with hygromycin (Hygro) or puromycin (Puro) resistance genes and GFP for their longitudinal tracking upon D-Luciferin (D-Luc) administration and selective removal with ganciclovir (GCV) or chemical inducer of dimerization (CID). (**B**) Bioluminescence signal normalized to wells with solvent of NSCs expressing HSV-TK, iC9, both (DS) or none, treated with 10 μM GCV for 6 days, 100 nM CID on day 0 and 5, or a combination of both (Comb) under proliferative conditions. (**C**) Cell counts on the last day of the treatment of safety switch NSCs under proliferation. (**D**) Representative confocal images of proliferative safety switch NSCs after 6 days of solvent (Ctrl) or Comb administration, stained with DAPI (grey) for the nucleus and expressing GFP (green). (**E**) Bioluminescence signal normalized to wells with solvent of rFluc or DS-NSCs, treated with 10 μM GCV for 6 days, 100 nM CID on day 0 and 5, or a combination of both under differentiation conditions. (**F**) Cell counts on the last day of the treatment of rFluc or DS-NSCs kept in total for 2 weeks under differentiation conditions. Data obtained from three independent experiments. Boxplots show the median (center line), IQR (box), and whiskers extending to 1.5 x IQR, with red diamonds indicating the mean. Ribbon plots display the mean ± SEM. Statistical significance was defined as *p < 0.05, **p < 0.01.

NSCs under proliferative conditions were treated with 10 μM GCV for 6 consecutive days, 100 nM CID on days 0 and 5 for 24 hours, or a combination of both (Comb). The bioluminescence signal decreased to 13.2 ± 6.8% and 11.2 ± 5.6% on the first day for iC9 treated with CID or Comb, respectively (**Fig. 1B**). Similar values were observed after 1 day for DS-NSCs, for which the bioluminescence signal was reduced to 15.3 ± 3.2% with CID and even lower to 12.5 ± 3.1% with Comb. NSCs expressing HSV-TK reached similar levels on day 6 of treatment with GCV (20.5 ± 5.1%) or Comb (19.9 ± 5.2%). At this time point, the relative bioluminescence signal of NSCs expressing iC9 treated with CID (14.9 ± 2.5%) or Comb (14.1 ± 2.1%) was slightly higher than on day one, probably due to the development of resistance. The bioluminescence signal of DS-NSCs decreased to 24.7 ± 7.6% with CID, to 20.9 ± 2.5% with GCV, and reached the lowest value with Comb (4.1 ±1.3%). We then recorded the cell count at the end of the experiment to ensure that the bioluminescence signal correlated with cell death (**Fig. 1C, D**). iC9 NSC counts were reduced to 36.3 ± 34.5% with CID and 29.1 ± 28.41% with Comb, which suggested that the killing efficiency was not as good as expected, given the bioluminescence results. On the contrary, HSV-TK NSC counts significantly decreased when treated with GCV (15 ± 7.2%, p = 0.0125) or Comb (12.6 ± 6.4%, p = 0.0107) compared to control. The reduction in cell count was even more pronounced in DS-NSCs treated with CID (12.3 ± 10.8%, p = 0.00735), GCV (5.93 ± 4%, p = 0.00484), and especially Comb (0.8 ± 0.7%, p = 0.0035).

We then carried out a similar experiment under differentiation conditions to better understand whether HSV-TK dependence on the cell cycle could affect the killing efficiency. NSCs expressing DS or only rFluc were kept for 8 days before the start of the treatment without small molecules or FGF to induce differentiation, and these conditions were maintained during the treatment period. After the first day of treatment, the bioluminescence signal of DS-NSCs decreased to 31.3 ± 4.3% with CID and to 32.3 ± 3.3% with Comb **(Fig. 1E**). Five days later, the relative bioluminescence signal of DS-NSCs treated with CID was not as low (43.9 ± 11.1%), and DS-NSCs treated with GCV reached similar values (52.5 ± 23.5%). Again, the Comb treatment was the most effective, resulting in a reduction of bioluminescence signal to 17.3 ± 4.3 % compared to the solvent control. Cell counts of DS-NSCs decreased when they were treated with CID to 60.5 ± 19.3% or with GCV to 50.1 ± 22.7% **(Fig. 1F**). However, the decrease in cell count was only significant when treated with Comb (24 ± 8.1%, p = 0.0194).

In conclusion, although individual switches have shown limited success, it is apparent that a combination treatment of CID and GCV is significantly superior at selectively removing DS-NSCs under proliferation and, to a lesser extent, differentiation conditions. Therefore, this safety switch system was chosen to further determine the effects of NSC removal in a stroke mouse model.

### Ablation of dual safety switch NSC in stroke-induced mice reduces graft size and proliferation rate

DS-NSCs were injected intracerebrally into immunodeficient mice 7 days after photothrombotic stroke induction.^21,22^ Two daily injections of solvent (Ctrl) or GCV 50 mg/Kg were administered for 2 weeks from day 15 post-stroke, combined with a daily injection of 10 mg/Kg CID for 5 days starting at day 15 and 25 post-stroke (Comb) (**Fig. 2A**). The bioluminescence signal in the head region was measured longitudinally (days 7, 15, 20, 25, 30, 35, 42 post-stroke) to determine the presence of DS-NSCs (**Fig. 2B**). Five days after the initiation of the treatment, a significantly lower bioluminescence signal was observed in the treatment group compared to control mice, (day 20: 16.7 ± 20.1%, p = 0.0001), and this difference remained until the end of the treatment (day 30: 17.2 ± 13.4%, p = 0.0002) (**Fig. 2C).** The normalized bioluminescence signal of the Ctrl group significantly increased almost ten-fold from the start of the treatment (5.1 x10^6^ ± 7.2 x10^6^) until the end of the experiment (48.3 x10^6^ ± 35 x10^6^, p = 0.0002) (**Fig. 2D**). On the contrary, the normalized bioluminescence signal of the Comb group was only slightly increased (from 3.2 x10^6^ ± 2.7 x10^6^ to 7 x10^6^ ± 5.3 x10^6^) by the end of the experiment and it was significantly lower than the Ctrl group (p < 0.0001).

**Figure 2.**
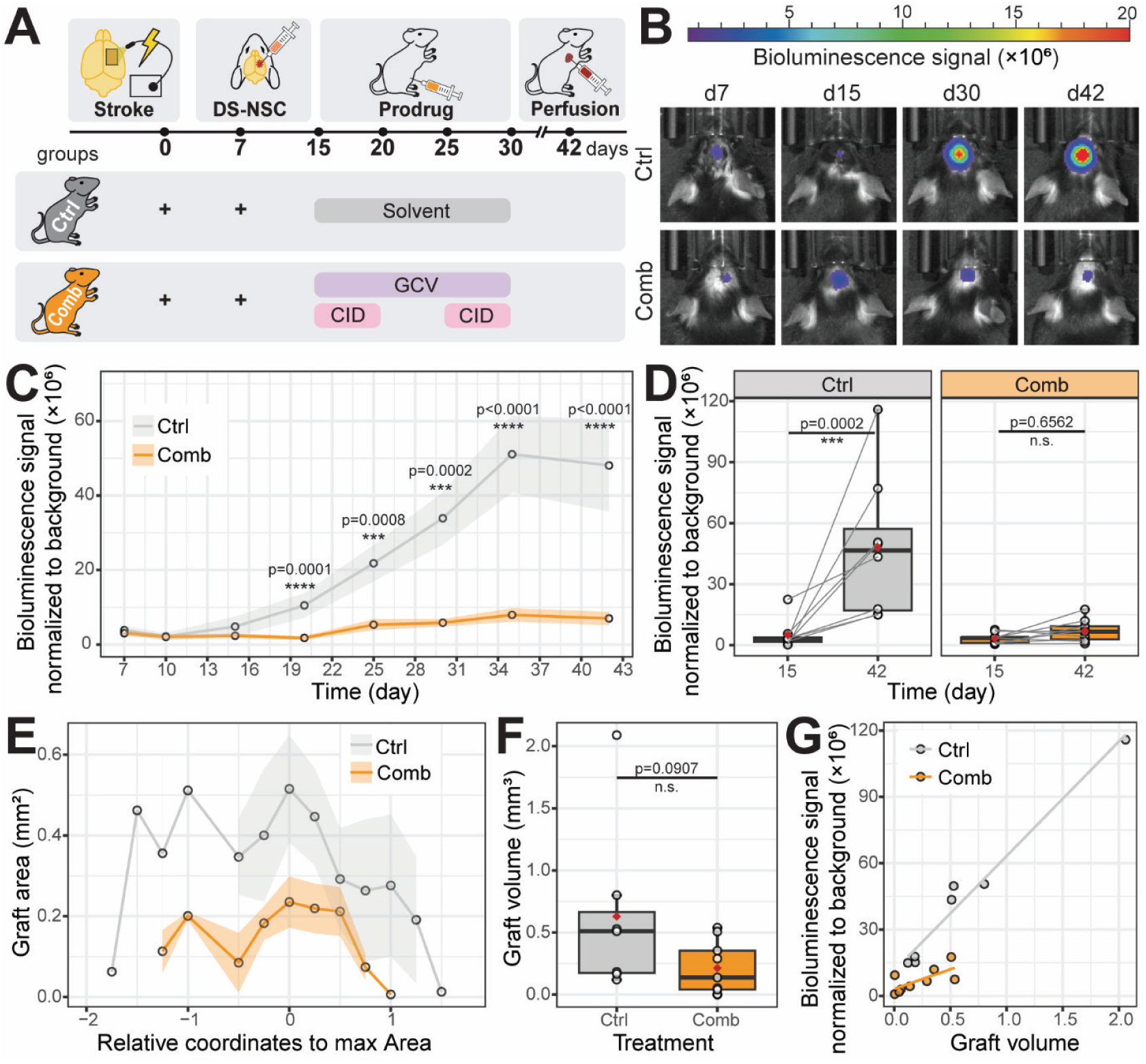
Dual switch NSCs are selectively removed following intracerebral injection in stroke mice. (**A**) Experimental design of DS-NSCs removal in a photothrombotic stroke mouse model. (**B**) Representative bioluminescence images of mice transplanted 7 days post-stroke with DS-NSCs and injected intraperitoneally with 50mg/Kg GCV twice a day for 2 weeks and 10 mg/Kg CID once a day for 5 days starting at day 15 and 25 (Comb) or with the solvent (Ctrl). (**C**) Bioluminescence signal from the head region of mice receiving DS-NSCs and treated with GCV and CID (Comb, orange) or the solvent (Ctrl, grey). (**D**) Bioluminescence signal paired data on the first day of the treatment (day 15) and on the last day of the experiment (day 42). (**E**) Graft area (mm^2^) of different brain sections centered on the slice with the maximum area of control (grey) and treated (orange) mice. (**F**) Graft volume (mm^3^) calculated by the trapezoidal integration of the adjacent area measurements. (**G**) Correlation between the bioluminescence signal on day 42 and the graft volume of Ctrl (grey) and Comb (orange) groups. Boxplots show the median (center line), IQR (box), and whiskers extending to 1.5 x IQR, with red diamonds indicating the mean. Ribbon plots display the mean ± SEM. Statistical significance was defined as n.s. p>0.05, ***p < 0.001, and **** p < 0.0001.

We measured the graft area across different brain sections to better characterize the presence of transplanted cells at the end of the treatment, and the area was consistently bigger in the control group (**Fig. 2E**). Accordingly, graft volume tended to be larger in the control group (0.63 ± 0.69 mm^3^) compared to the treatment group (0.21 ± 0.22 mm^3^) (**Fig. 2F**). We then correlated the bioluminescence signal on day 42 to the graft volume and observed a positive correlation in both groups (**Fig. 2G**). Nevertheless, the control group had a stronger linear correlation (R^2^= 0.98) than the Comb group (R^2^= 0.49), and the slope was almost 3 times higher in the control group (51.8 x10^6^ vs.17.7 x10^6^). Taken together, these results showed that the treatment group had fewer NSCs expressing firefly luciferase on the last day of treatment than the control group.

To analyze the graft proliferation rate, we injected EdU on day 15 or 20 and BrdU on day 25 or 30 post-stroke, and we further stained for Ki67 to label dividing cells at the end of the experiment (**Fig. 3A, 3F**). In general, the number of positive cells in both groups decreased over time (EdU > BrdU > Ki67). EdU injected at the onset of the treatment (day 15) revealed a significantly lower number of labeled cells in the treatment group at the end of the experiment (25.1 ± 15 vs. 63.4 ± 25.2, p = 0.0192), consistent with the elimination of proliferating NSCs once GCV and CID were administered (**Fig. 3B**). Five days following GCV and CID administration (day 20), the mean count of EdU^+^ cells was again significantly smaller in mice with the combination treatment (31.4 ± 21.4 vs. 98.3 ± 41.3, p = 0.0357). Interestingly, the number of BrdU^+^ NSCs at day 25 tended to be higher in the treatment group (7.5 ± 2.2 vs. 6.3 ± 2.6). However, the tendency changed again on day 30 (Ctrl: 5.4 ± 2.7 vs. Comb: 2.6 ± 1.8) (**Fig. 3C**). At the end of the experiment (day 42), the number of Ki67^+^ cells was significantly higher in the control group (1.6 ± 0.6) compared to mice receiving the combination treatment (0.4 ± 0.6, p = 0.00391) (**Fig. 3D**). Consistent with this reduced proliferation, the total number of human nuclei (HuNu^+^) in the field of view was also significantly lower in the mice receiving the treatment (367 ± 218 vs. 630 ± 81.1, p = 0.00587) (**Fig. 3E**).

**Figure 3.**
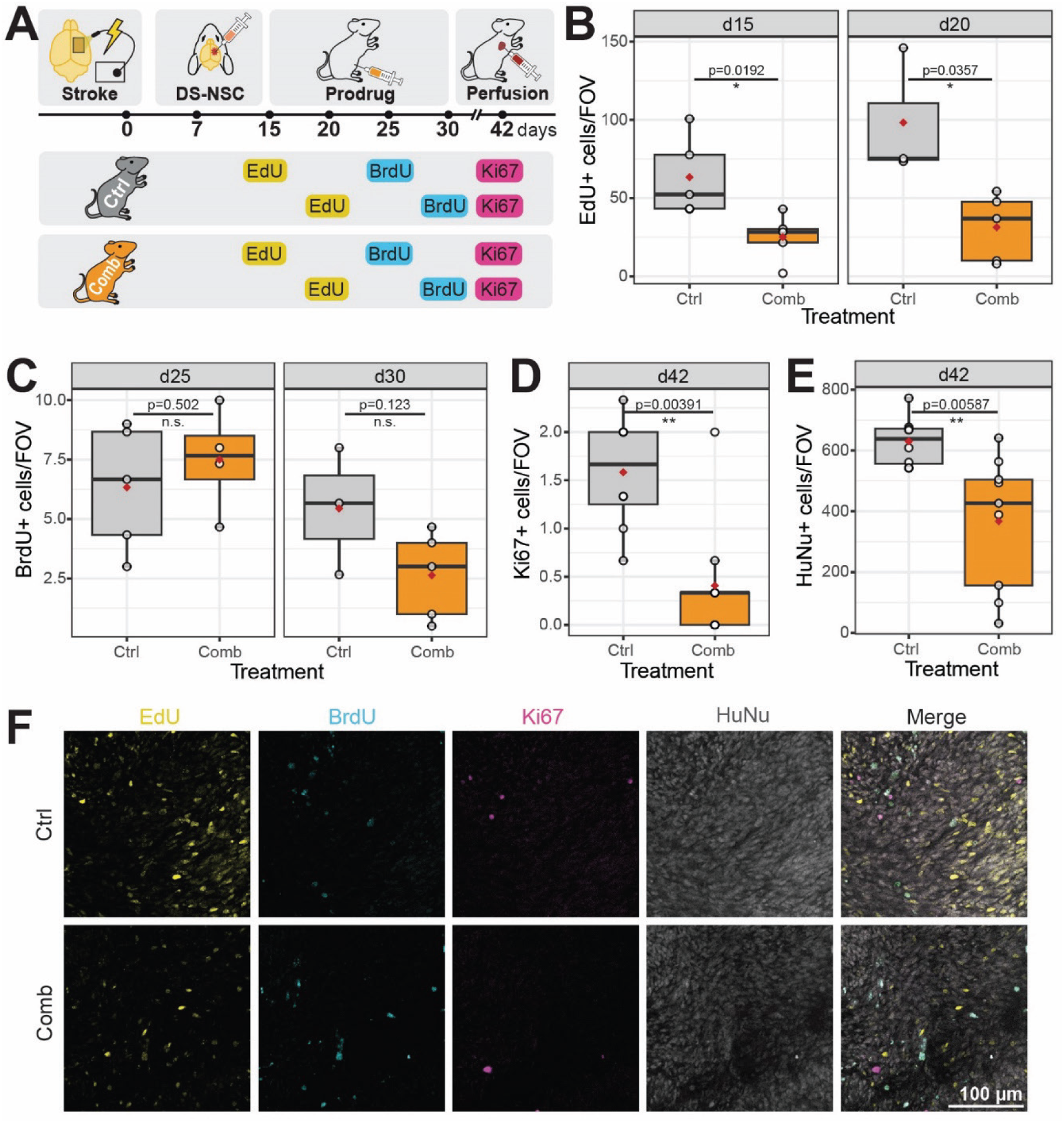
Lower proliferation rate of dual switch NSCs in mice receiving GCV and CID. (**A**) Experimental design of nucleoside analogue injection for determining the proliferation of DS-NSCs. (**B**) Number of EdU^+^ cells per field of view (FOV) of mice receiving EdU on day 15 (left) or 20 (right) post-stroke. (**C**) Number of BrdU^+^ cells per FOV of mice receiving BrdU on day 25 (left) or 30 (right) post-stroke. (**D**) Number of Ki67^+^ or (**E**) human nucleus (HuNu^+^) cells per FOV of mice on the last day of the experiment (day 42). (**F**) Representative 63x confocal images of mice transplanted with DS-NSCs and receiving EdU (yellow) at day 15, BrdU (turquoise) at day 25, and stained for Ki67 (magenta) and HuNu (white) at the end of the experiment. Boxplots show the median (center line), IQR (box), and whiskers extending to 1.5 x IQR, with red diamonds indicating the mean. Statistical significance was defined as n.s. p>0.05, *p < 0.05, **p < 0.01.

In conclusion, mice getting the combination treatment presented fewer NSCs and a lower proliferation rate compared to the control group. Therefore, we carried out histological and behavioral analysis to further understand whether the death of the grafted NSCs affects the stroke recovery.

### Dual safety switch NSC removal leads to larger, more inflamed, and poorly revascularized stroke lesions

Mouse brain slices were stained with antibodies targeting glial fibrillary acidic protein (GFAP) and ionized calcium-binding adaptor molecule 1 (IBA1) to delimitate and characterize the stroke lesion, and in addition, DAPI and GFP were used to define the graft boundaries (**Fig. 4A**). Comb-treated mice consistently exhibited larger stroke areas across different brain sections (**Fig. 4B**). In consequence, the stroke volume also tended to be larger in the Comb group (0.95 ± 0.43 mm^3^) than the control mice (0.73 ± 0.3 mm^3^) (**Fig. 4C**). To further understand how the removal of NSCs influences the inflammatory status of the stroke lesion, we analyzed the fluorescence intensity of GFAP and IBA1 in the ischemic border zone (IBZ) (**Fig. 4D, E**). The mean normalized GFAP fluorescence intensity was similar in both groups (Comb: 6.37 ± 1.89 vs. Ctrl: 5.36 ± 1.4) (**Fig. 4F**). However, the mean normalized IBA1 fluorescence signal was significantly higher in the IBZ of treated mice (2.31 ± 0.35) compared to vehicle-treated mice (1.91 ± 0.28, p=0.0211) (**Fig. 4G**).

**Figure 4.**
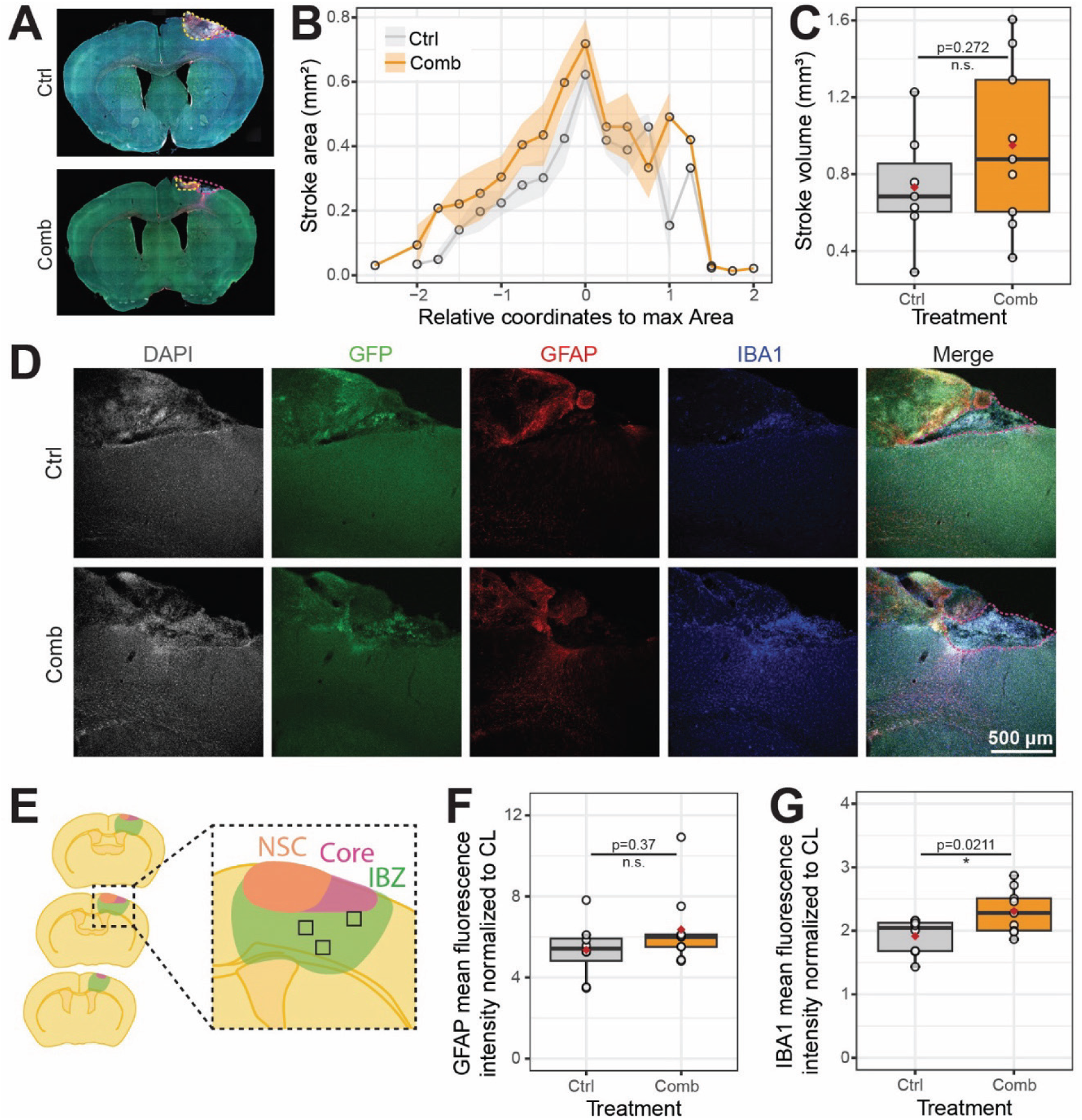
Dual switch NSC removal from the stroke lesion leads to larger stroke lesions exhibiting a pro-inflammatory state. (**A**) Representative slide scanner images depicting the stroke lesion (pink dashed line) and the graft (yellow dashed line). (**B**) Stroke area (mm^2^) of different brain sections centered on the slice with the maximum area of control (grey) and treated (orange) mice. (**C**) Stroke volume (mm^3^) calculated by the trapezoidal integration of the adjacent area measurements. (**D**) Representative 10x confocal images of brain slices containing GFP^+^ (green) cells stained for DAPI (white), GFAP (red), and IBA1 (blue). (**E**) Schematic representation of the NSC graft (orange), the stroke core region (pink), and the ischemic border zone (IBZ, green). (**F**) Mean of GFAP fluorescence signal measured in three ROIs in the IBZ of 5 brain slices and normalized to the contralateral (CL) hemisphere of control (grey) and treated mice (orange). (**G**) Mean of IBA1 fluorescence signal measured in three ROIs in the IBZ of 5 brain slices and normalized to the contralateral (CL) hemisphere of control (grey) and treated mice (orange). Boxplots show the median (center line), IQR (box), and whiskers extending to 1.5 x IQR, with red diamonds indicating the mean. Ribbon plots display the mean ± SEM. Statistical significance was defined as n.s. p>0.05 and *p < 0.05.

To measure the vascularization of the brain, laser Doppler imaging (LDI) was carried out directly after the stroke induction and at the end of the experiment (**Fig. 5A**). In both groups, the stroke induction resulted in approximately a 50% decrease of the relative blood perfusion in the right hemisphere normalized to the contralateral side (Ctrl: 49.2 ± 12.3%, Comb: 46.8 ± 9.3%) (**Fig. 5B, C**). All mice had significantly higher blood perfusion at the end of the experiment compared to the day of the induction (p < 0.0001). Nevertheless, on day 42, the mean relative blood perfusion of mice in the control group tended to be higher (83.8 ± 14.4%) than that of the treatment group (74.7 ± 10.9%).

**Figure 5.**
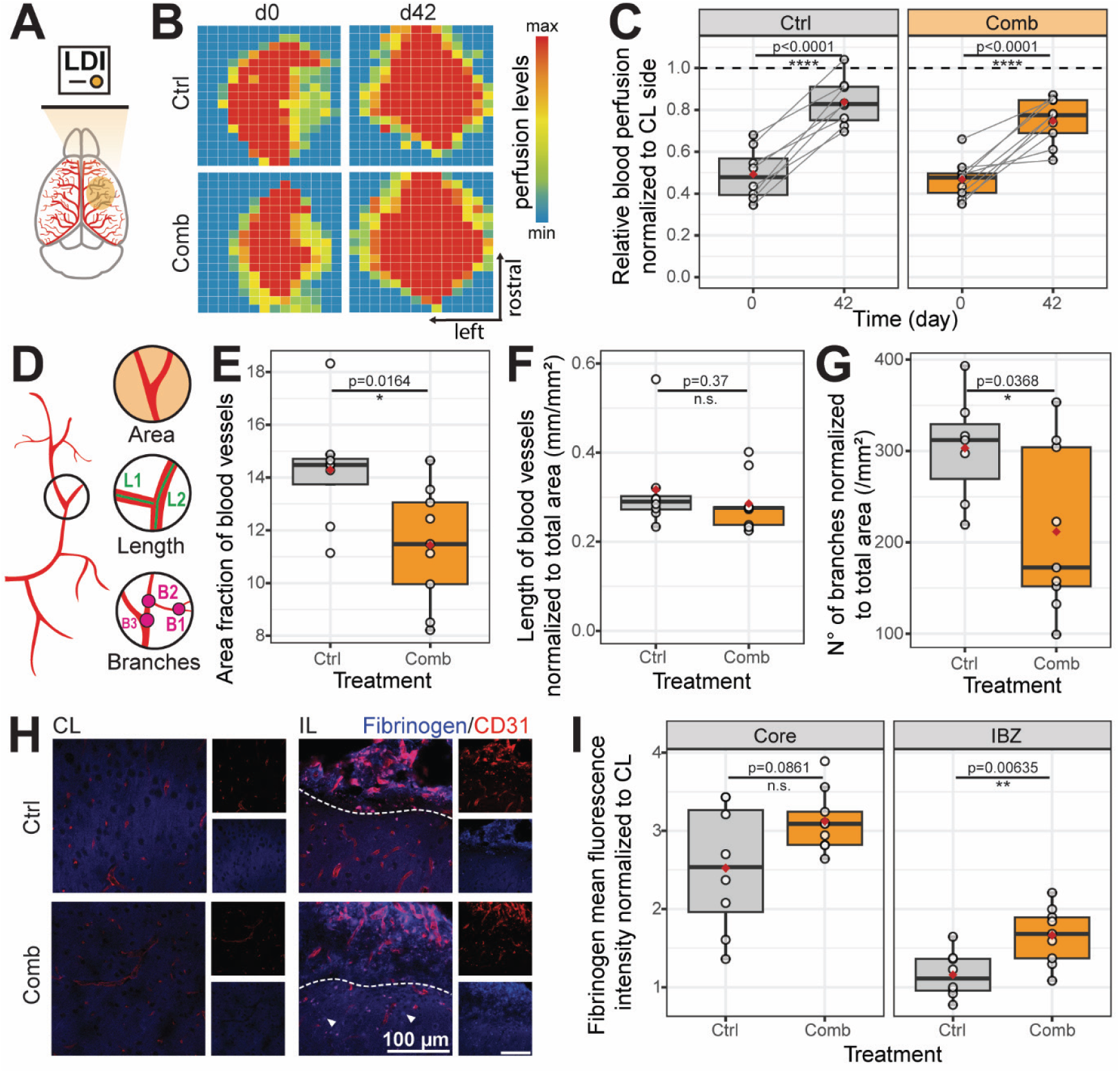
Ablation of dual switch NSCs results in poor revascularization. (**A**) Schematic of Laser Doppler Imaging (LDI) (**B**) Representative images of LDI after stroke induction (day 0) and before the end of the experiment (day 42). (**C**) Paired data of relative blood perfusion measured with LDI normalized to the contralateral (CL) hemisphere. (**D**) Schematic of the analyzed vasculature characteristics. (**E**) Area, (**F**) length, and (**G**) number of branches of blood vessels, normalized to the selected ROI area in the ischemic border zone (IBZ). (**H**) Representative 63x confocal images of brain slices stained with CD31 (red) and fibrinogen (blue) antibodies used for the vasculature analysis of the contralateral (CL) and ipsilateral (IL) hemispheres. (**I**) Mean of fibrinogen fluorescence signal measured in three ROIs outside blood vessels in the core (left) and IBZ (right) of 3 brain slices and normalized to the contralateral (CL) hemisphere of control (grey) and treated (orange) mice. Boxplots show the median (center line), IQR (box), and whiskers extending to 1.5 x IQR, with red diamonds indicating the mean. Statistical significance was defined as n.s. p>0.05, *p < 0.05, **p < 0.01, and **** p < 0.0001.

To further understand the vascular changes associated with NSC removal, we carried out an immunostaining with antibodies against CD31 for characterizing blood vessel area, length, and number of branches (**Fig. 5D**). The area fraction of blood vessels was significantly smaller in mice receiving the combination treatment (11.4 ± 2.2) than in control mice (14.3 ± 2.1, p = 0.0164) (**Fig. 5E**). The blood vessel length normalized to the vascular area was similar between both groups (Comb: 0.29 ± 0.06 mm/mm^2^, Ctrl: 0.32 ± 0.1 mm/mm^2^) (**Fig. 5F**). Besides, the treatment group had significantly fewer branches normalized to the total vascular area (211.8 ± 90.7) than the control group (303.2 ± 58.7, p = 0.0368) (**Fig. 5G**). We then performed immunostaining of fibrinogen to analyze the permeability of the newly formed blood vessels (**Fig. 5H**). The mean fibrinogen fluorescence intensity outside blood vessels tended to be higher in the core region of treated mice (3.12 ± 0.39 vs. 2.52 ± 0.81), and it was significantly higher in the IBZ of the Comb group (1.66 ± 0.36) compared to control (1.16 ± 0.29, p = 0.00635) (**Fig. 5I**).

Together, these results show that despite the ablation efficiency, the removal of DS-NSCs led to bigger stroke lesions with a more pro-inflammatory environment, and an impaired revascularization.

### NSC ablation impairs contralateral hindlimb motor recovery after stroke

To evaluate the functional effects of NSC removal, mice were tested repeatedly on a horizontal ladder walk, and hindlimb placement accuracy was analyzed using the deep learning-based tool DeepLabCut (**Fig. 6A**), as previously described.^23^ Error rate of the left hindlimb, contralateral to the photothrombotic stroke, was used as the primary behavioral outcome. Prior to stroke induction, both groups displayed comparably low error rates (Comb: 2.00 ± 1.10%; Ctrl: 1.22 ± 0.67%; p > 0.05), confirming no pre-existing motor differences (**Fig. 6B**). On day 3 post-stroke, error rates increased markedly in both groups to similar levels (Comb: 26.67 ± 5.16%; Ctrl: 22.73 ± 4.67%; p > 0.05), indicating equivalent initial stroke-induced motor deficits. No significant differences between groups were detected on days 7, 14, or 21. However, from day 28 onward, the groups significantly diverged. Mice retaining the NSC graft (Ctrl) showed progressive recovery toward pre-stroke levels, whereas mice in which NSCs were ablated (Comb) maintained elevated error rates. Pairwise comparisons using estimated marginal means confirmed significant differences at day 28 (p < 0.001), day 35 (p = 0.041), and day 42 (p = 0.049). Importantly, no group differences were observed for the ipsilateral hindlimb (right back toe tip) at any time point (all p > 0.05), consistent with unilateral stroke confined to the right hemisphere.

**Figure 6.**
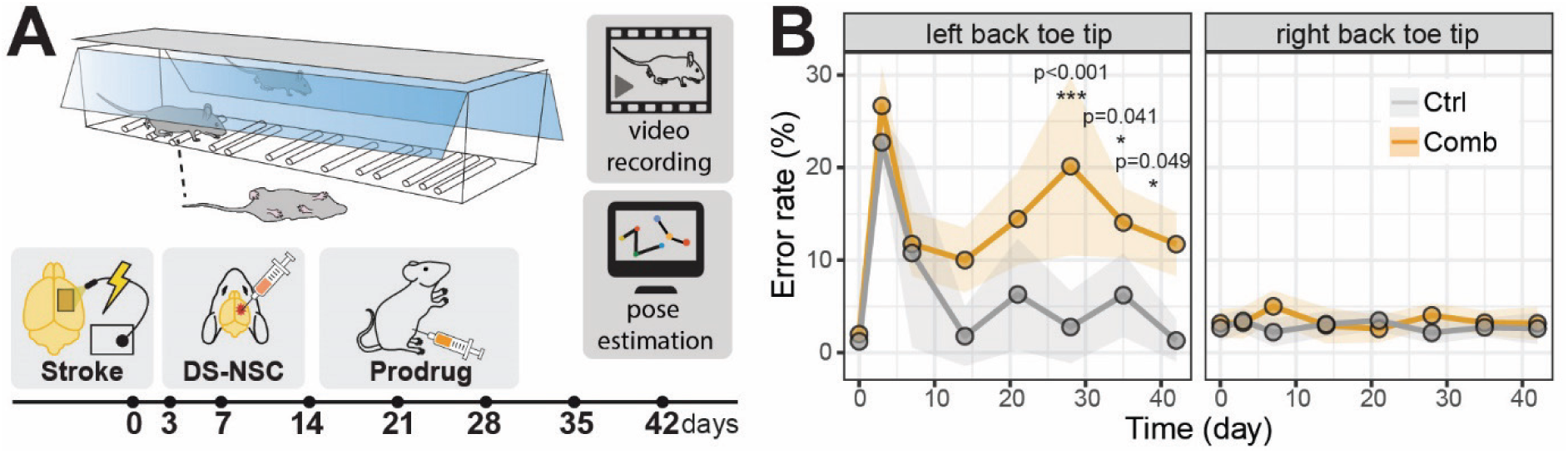
NSC ablation impairs contralateral hindlimb motor recovery after stroke. (**A**) Schematic of the horizontal ladder walk test setup. Mice were recorded at days 0, 7, 14, 21, 28, 35, and 42 post-stroke, and limb placement was analyzed using DeepLabCut pose estimation. (**B**) Error rate (%) of left back toe tip (contralateral to stroke, left panel) and right back toe tip (ipsilateral, right panel) over time in control (grey, n = 8) and combination treatment (orange, n = 9) groups. Estimated marginal means from a linear mixed-effects model. Data presented as mean ± SEM. *p < 0.05, ***p < 0.001.

In sum, these data indicate that the physiological consequences of DS-NSCs ablation described above are accompanied by impaired motor recovery in the contralateral hindlimb, whereas graft retention promotes functional improvement.

## Discussion

In this study, we developed and validated a dual safety switch system combining HSV-TK and iC9 for the selective elimination of NSC grafts in a murine stroke model. Our findings demonstrate that while the dual safety switch enables graft removal, it comes at the cost of limiting regeneration-associated tissue responses and compromising motor recovery.

NSCs under proliferation transduced with lentiviral vectors containing HSV-TK or iC9 are selectively removed upon GCV or CID administration, respectively. Previous groups have reported that iC9 is faster, more efficient, and safer than HSV-TK,^24,25^ which aligns with our bioluminescence results. Nevertheless, no significant differences were found in the cell count of iC9 NSCs on the last day of the treatment. Considering that the mean number of iC9 NSCs treated with solvent tended to be lower compared to the other cell lines, we suggest that NSCs, containing very high levels of iC9, dimerize this monomer in the absence of CID triggering a premature apoptosis. Nevertheless, as previously reported, the initial expression of iC9 highly influences the killing efficiency,^18^ and thus, isolating NSCs that present low levels of iC9 might result in death escape.

Therefore, to enhance the killing efficiency, NSCs were transduced with both safety switch systems. We demonstrated that bioluminescence signal and cell count decreases more than 95% when treated with a combination of GCV and CID, significantly surpassing the death rate of single safety switches. This increase in killing efficiency, nevertheless, comes along with a higher risk of insertional mutagenesis, which could be prevented by targeting safe harbor sites such as the adeno-associated virus integration site 1 (AAVS1), gene trap ROSA 26, or citrate lyase beta-like (CLYBL).^26^ Another strategy consists of generating a transgene-free safety switch system by disrupting the uridine monophosphate synthetase gene, which renders the proliferation dependent on external uridine.^27^

The dual switch developed in this study targets both proliferating and differentiating NSCs. However, when NSCs were kept under neural differentiation conditions for 8 days before the initiation of the treatment, the killing efficiency was decreased compared to proliferative conditions. *In vitro* differentiation results resemble *in vivo* outcomes, since the grafted cells were not completely removed. The correlation slope between bioluminescence signal and graft volume differs between treatment groups, indicating that for a given graft volume, the Comb group exhibited a lower bioluminescence signal. There are several possibilities for such readout: incomplete revascularization hinders D-luciferin delivery to the graft, transgenes (including rFluc) were silenced, or safety switch cells were ablated leading to the positive selection of non-transduced NSCs. Indeed, it has been described that although EF1a is a strong promoter and efficient for NSCs,^28^ it can be silenced through the differentiation process due to DNA methylation at CpG islands.^29^ Therefore, the choice of the promoter is crucial for ensuring a stable suicide gene expression. Promoters can target specific populations, e.g., pluripotent undifferentiated cells by using NANOG or human OCT4 short response element (hOCT4-SRE), or exclude certain populations, such as differentiated neurons, via Ki67.^30,15^ The expression can be temporarily regulated via the Tet-inducible system for reducing metabolic side effects of HSV-TK, or it can be ubiquitous, such as the actin beta (ACTB) promoter, to ensure high and stable levels.^31^ Another approach to overcome gene silencing is to carry out biallelic integration into the AAVS1 safe harbor locus.^32^

To further characterize the graft proliferation, we injected nucleoside analogues at regular time intervals, and the treatment group had persistently fewer dividing cells. The reduction in the proliferation rate achieved during GCV and CID injection was maintained after the termination of the treatment, which is crucial for preventing relapses. This result is limited to five time points and thus, to further understand the proliferation of grafted cells in the presence and absence of the treatment, it would be interesting to use a more complex system, such as the “Brainbow”, which is based on Cre/lox stochastic recombination of different fluorophores, enabling the tracking of the cell progeny.^33^

Regarding the inflammatory status, we found that mice receiving the combination treatment 8 days after transplantation showed higher IBA1 fluorescence signal levels in the IBZ. The recruitment of microglia and/or infiltrating macrophages to the IBZ of treated mice may result from the removal of transduced cells or from the absence of therapeutic NSCs. We had previously observed similar outcomes in vehicle-treated compared to NSC-treated mice (higher levels of IBA1 in the ipsilateral hemisphere but no big differences in GFAP).^34^ Taking this together with the fact that apoptotic MSCs produce anti-inflammatory effects,^35^ we suggest that the pro-inflammatory status is due to the lack of grafted cells rather than the release of death-related factors, but it should be further validated with an orthogonal ablation system. Similarly, we demonstrated that the removal of NSCs has a negative impact on the revascularization process, leading to less dense blood vessels with fewer branches and higher permeability. To ensure xeno-graft survival, we injected human NSCs into an immunodeficient mouse model (Rag2-/-) that was previously described to differ in inflammation and vascular repair from immunocompetent stroke mouse models.^36^ Therefore, it would be crucial to repeat these experiments in immunocompetent mice with allogenic mouse stem cells or in humanized mouse models to ensure the clinical translation. Even so, the incomplete revascularization observed in this study could be a consequence of the lack of sustained growth factors, suggesting that the maintenance of grafted cells is crucial for the complete recovery of stroke symptoms. In this line, the use of biomaterials that mimic the extracellular matrix, such as hydrogels, could be used to release growth factors in a sustained and adjustable manner,^37–39^ which could further decrease the inflammation, improve the revascularization, and increase the survival of transplanted stem cells.

Histological findings correlate with physiological outcomes. Mice receiving solvent injections restored the hindlimb error rate to baseline timepoint, whereas the treated mice showed a sustained error rate from day 28 until the end of the experiment. These results suggest that the NSC effect goes beyond the initial supply of beneficial factors. Nevertheless, further experiments need to be conducted to understand how the circuitry is affected by the ablation of transplanted stem cells, via, e.g., monosynaptic rabies virus tracing.^40^

Overall, our findings demonstrate that DS-NSC removal from the stroke lesion is feasible and helps control graft volume and proliferation, but this comes at the expense of the recovery process. The lack of complete anatomical and functional restoration following graft ablation suggests that the sustained presence of stem cells is indispensable for driving comprehensive repair. Consequently, these results reinforce the clinical value of stem cell grafts over their transient derivatives for neurological disorders.

## Methods

### Experimental design

This study was designed to analyze the effect of human iPSC-derived NSC removal in the stroke pathology of immunodeficient *Rag2*^-/-^ mice. We hypothesize that the administration of GCV and CID 1 week after transplantation of DS-NSCs results in (1) selective removal of grafted cells, (2) lower recovery-associated tissue responses, and (3) worse recovery in motor function compared to the control group. To test this, mice underwent photothrombotic stroke induction followed by intracerebral NSC injection, and they were allocated into two groups: (1) treatment group (*n* = 10) received the combination treatment of GCV and CID, (2) the control group (*n* = 10) received only the solvent, serving as a negative control. Photothrombotic stroke induction was validated by performing laser-Doppler imaging (LDI) 30 min after stroke onset, and additionally before perfusion (on day 42). The bioluminescence signal was measured at regular intervals (every 5 days during treatment administration and every 7 days afterwards). Motor function was assessed weekly by performing the horizontal ladder walk test, which was further analyzed with DeepLabCut (DLC). After perfusion, histological analysis was performed to characterize the presence and proliferation of grafted cells and evaluate stroke pathological markers. Animals that did not present a bioluminescence signal at any time point, indicating unsuccessful intracerebral injection (regardless of their group) were excluded (Ctrl *n* = 8, Comb *n* = 9). Another mouse was excluded from the control group for the calculations of the graft and stroke areas and volumes, as the graft was accidentally removed during perfusion.

### Animals

All animal experiments were performed at the Laboratory Animal Services Center (LASC) in Schlieren according to the local guidelines for animal experiments and were approved by the Veterinary Office in Zurich, Switzerland (License ZH110/2023). In total, 20 adult male and female genetically immunodeficient *Rag2*^-/-^ mice (weight range: 20-32 g) were employed for this study. Mice were kept in top-filter laboratory cages under OHB conditions in a temperature and humidity-controlled room with a constant 12/12 h light/dark cycle (light on from 6:00 a.m. until 6:00 p.m.). All mice were standard housed in groups of at least two and maximal four per cage with ad libitum access to standard diet food pellets and water.

### Cell culture

We previously established a xeno-free differentiation protocol of human iPSCs to NSCs.^21,28,41^ NSCs were maintained with Neural Stem cell Maintenance Medium (NSMM) (50% DMEM/F12, 50% Neurobasal medium, 1x N2 supplement, 1x B27 supplement, 1x Glutamax) supplemented with small molecules (10 ng/ml hLIF, 4 μM CHIR99021, 3 μM SB431542) and 5 ng/mL FGF. For the spontaneous NSC differentiation, NSCs were maintained with NSMM without small molecules or FGF for a total of 2 weeks. Transduced NSCs were treated with 0.5 μg/mL of puromycin and/or 0.2 mg/mL of hygromycin daily up to the beginning of the treatment.

### Cloning

The dual-reporter lentiviral construct pLL410_EF1a-rFLuc-T2A-GFP-mPGK-Puro (LL410PA-1) obtained from System Bioscience was used as the starting backbone. HSV-TK and iC9 sequences were amplified with KAPA HiFi hot start mix (Roche, 07958927001) followed by a PCR purification (Sigma, NA1020-1KT). After digestion with BstBI (New England Biolabs, R0519S) and XbaI (New England Biolabs, R0145S) of the plasmid and inserts, the desired DNA fragments were purified from an agarose gel (Sigma, NA1111). Quick ligase (Biolab, M2200S) was used to ligate the inserts into the plasmid in a 3:1 molar ratio for subsequent transformation of NEB® 5-alpha F’Iq Competent E. coli (Biolabs, C2992H). We carried out a PCR in 9 colonies per construct with KAPA2G Fast Genotyping (KAPA Biosystems, KK56121) and incubated them in LB media for 6 h at 37 °C with agitation at 1000 rpm. Two positive colonies were further sequenced to ensure the lack of mutations before proceeding with the GenElute™ Endotoxin-free Plasmid Maxiprep Kit (Sigma Aldrich, PLEX15-1KT). A similar protocol using BstBI (New England Biolabs, R0519S) and BstEII-HF (New England Biolabs, R3162S) restriction enzymes was followed to exchange GFP, mPGK promoter, and puromycin resistance gene with the hygromycin resistance gene.

### Lentiviral production

We conducted the calcium phosphate transfection as previously described.^42^ In brief, packaging plasmids and transfer vector were mixed with water containing HEPES, CaCl_2,_ and 2x HBS. The mix was added to LentiX HEK cells, and the media (DMEM + 10% FCS) was changed the day after. For iC9 lentiviral production, LentiX HEK cells were supplemented with Pan Caspase OPH Inhibitor Q-VD (R&D, OPH001-01M) to prevent the premature apoptosis of packaging cells. Two days later, virus was harvested and ultracentrifuged (25,000 x g, 4 °C, 1 h 30 min) before final resuspension in MEM and snap-freezing in liquid nitrogen.

### Virus transduction and titration

A serial dilution of virus concentrate (0, 1, 2, 4, 8, 16 μL) was added directly to 6 wells of a 24-well plate containing 120,000 NSCs/well. Virus titer was determined by FACS for HSV-TK or survival rate to hygromycin for iC9. The selected volume was upscaled to transduce 600,000 NSCs/well in a 6-well plate, and after 24 h, antibiotics were added to select transduced NSCs. In addition, HSV-TK NSCs were sorted to select a population with high expression of GFP.

### Treatment administration

GCV (MedChemExpress, HY-13637) was diluted freshly with the solvent mix (4% EtOH, 10% PEG-400, 1.7% Tween-20 in water) at a concentration of 5 mg/mL, followed by sonication and a 5-minute incubation in a shaker at 65 °C and 1000 rpm. CID (TargetMol, T14299) was diluted to 62.5 mg/mL in EtOH, aliquoted, and stored at −20 °C. *In vitro*, NSCs were treated with 10 μM GCV for 6 days and/or 100 nM CID for 24 h at day 0 and 5 of the experiment. For *in vivo* experiments, GCV was injected i.p. twice a day at a 50 mg/kg dose for 14 consecutive days. CID was further diluted in the solvent mix containing GCV to reach a 1 mg/mL concentration and used within 30 min following dilution. CID was injected i.p. daily for 5 days starting on days 15 and 25 at a 10 mg/Kg dose.

### Bioluminescence imaging

For *in vitro* experiments, 60,000 NSCs/well were seeded in three wells of a 24-well plate for each treatment and cell line. The bioluminescence signal was measured on days 0, 1, 3, and 6 with a plate reader (Tecan M1000 pro). Bioluminescence signal of the well was measured as background for further normalization. The stock solution of 30 mg/mL D-Luciferin (Merk, 50227) was diluted in NSMM+SM media to reach a final concentration of 150 µg/mL, which was incubated with NSCs for 10 min at 37 °C before measuring the bioluminescence signal again.

For *in vivo* experiments, the head region of the mice was shaved with an electric razor to achieve optimal signal. A solution of 30 mg/mL D-Luciferin in PBS was injected i.p. at a 300 mg/Kg dose. Images were acquired 10, 15, and 20 min after substrate injection with the IVIS^®^ Lumina III In Vivo Imaging System. Bioluminescence was analyzed with the *Living Image* software (v. 4.7.3) by setting a circular region of interest (ROI) (diameter = 1.8 cm) and placing it in the head region, the thorax as background signal, and an additional randomly selected region to determine the noise of the image. Signal was quantified using total photon flux (photons per second) and normalized to the noise and background signals. Data was collected in Microsoft Excel and plotted with R software as mean ± SEM (v. 4.3.1).

### Cell counting

On the last day of the treatment, cultured NSCs were detached with Accutase solution (Sigma, A6964-100ML) and centrifuged for 5 min at 300 x g. NSCs were resuspended in 0.5 mL and diluted 1:10 to determine automatically the number of alive cells with the Vi-CELL XR cell viability analyzer (Beckmann Coulter).

### Photothrombotic stroke induction

Photothrombotic stroke was induced as previously described.^43–45^ In brief, after anesthesia was induced with 4.5% isoflurane vaporized in oxygen, mice were placed into a stereotactic frame (Davids Kopf Instruments) for surgery. Isoflurane was maintained at 1-2% for the whole surgery, and temperature was kept by placing a heating pad underneath the mouse. After checking the reflexes, ear bars were gently inserted into the ears underneath the ear bone to fixate the skull. An incision was made along the longitudinal fissure, skin was retracted, and fascia was removed. A solution of 15 mg/ml Rose Bengal dissolved in NaCl was injected at a dose of 10 μl/g i.p. 5 min before illumination. An Olympus KL1500LCD (“4’’) cold light source illumination lamp [150W, 3010K] with a surface area of 4 mm x 3 mm was positioned on the right to bregma (−2.5 mm to + 2.5 mm medial/lateral and 0 mm to + 3 mm from Bregma) and the lighting was maintained for 10 min. The mouse was then placed in an empty cage to recover before LDI.

### Laser Doppler imaging (LDI)

Mice were placed under anaesthesia, and the skull was exposed through a midline skin incision, before subjecting them to LDI (Moors Instruments, MOORLDI2-IR). It was carried out after stroke induction and on the last day of the experiment, followed by stitching and post-op care or perfusion, respectively. LDI data were exported and quantified in terms of flux in a defined ROI using Fiji (v. 1.54p) and subsequently analyzed with R software (v. 4.3.1).

### NSC intracerebral injection

Transplantation of NSCs was conducted as described in previous studies.^46^ Briefly, mouse preparation was the same as for stroke induction up to the dye injection. We then set the injection coordinates [AP: + 0.5 / ML: + 1.5 / DV: - 0.8 (mm relative to bregma)] and drilled a small hole (0.8 mm diameter) through the skull. We drew 3 μl of cell suspension (10^5^ cells /μl) using a 26G needle into a 10 μL Hamilton syringe and changed the tip to a 35G needle for injection. The needle was guided automatically to the calculated depth at a slow, steady rate. The needle was further inserted 0.1mm and then retracted to the described coordinates to create a pocket. Cells were injected at a constant rate (5nl/s), and the needle was left in place for 4 min to avoid spillovers. Once the needle was retracted, Histoacryl^®^ was applied to seal the cavity and closed the wound, followed by standard post-op care.

### Nucleoside analogue injection

BrdU and EdU suspensions were prepared freshly on the day of the injection. EdU was diluted to 5 mg/mL in PBS and injected i.p. at a 50 mg/Kg dose on days 15 or 20 post-stroke induction. BrdU was diluted to 10 mg/mL in PBS and injected i.p. at a 100 mg/Kg dose on days 25 or 30 post-stroke induction.

### Tissue processing and immunostaining

Animals were perfused transcardially using Ringer solution for 2 min, followed by 4% PFA for 5 min. Brains were extracted and incubated for 8h in 4% PFA. After incubating for 3 days in a 30% sucrose solution, brains were cut into 40 μm coronal sections using a Thermo Scientific HM 450 microtome. Brain sections were washed with PBS and incubated with 500 μl blocking buffer consisting of donkey serum (5%) diluted in PBS + 0.1% Triton® X-100 for 1h at RT. Sections were then incubated with primary antibodies (**Suppl. Table 1**) on a shaker at approximately 90 rpm overnight at 4 °C. The following day, sections were washed with PBS and incubated with corresponding secondary antibodies (**Suppl. Table 2**) for 3 h at RT. FlexAble kits were prepared according to the manufacturer’s instructions and added directly to the secondary antibody solution. For EdU staining, brain sections were incubated for 30 min with the reaction mix buffer (1 x per well equates to 325 μl ddH_2_O, 25 μl 2M Tris, 100 μl 10mM CuSo4, 0.1 μl 10mM AlexaFluor 647, 100 μl 500 mM Ascorbic acid). Brain slices were additionally incubated for 15 min with 20 ng/µL DAPI (Sigma, D9542-1MG diluted 1:50 in PBS). Sections were then mounted onto Superfrost PlusTM microscope using Mowiol® mounting solution.

### Imaging and analysis

For graft and stroke size analysis, brain slices stained with HuNu or GFAP and IBA1, respectively, were imaged on the Zeiss Axio Scan.Z1 slide scanner. Fiji was used to re-scale the image, enhance the contrast, and manually draw a polygon surrounding the graft or stroke area. Brain slices were then assigned to stereotaxic coordinates (distance relative to Bregma) using a mouse brain atlas.^47^ We used a trapezoidal integration of the cross-sectional areas (*A*) to estimate the graft and stroke volumes with R software.^44,48^ The volume (*V*) was calculated by summing the areas of adjacent slices, defined by the distance (*Δx*) along the anterior-posterior axis:

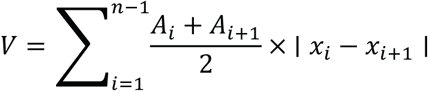

Brain sections were also imaged with a Leica SP8 laser confocal microscope with the 63x objective (for cell counting and permeability study) or the 10x objective (for inflammation and vasculature analysis). We customized a JavaScript to semi-automatically count HuNu^+^ cells with Fiji, and the ‘cell counter’ plugin was used to manually count EdU^+^, BrdU^+^, and Ki67^+^ cells. To analyze the inflammatory status, three rectangular ROIs (60×30 pixels) were placed in the ischemic border zone (IBZ) adjacent to the stroke core and the contralateral side for normalization. We analyzed five brain slices per mouse and calculated the mean fluorescence intensity of GFAP and IBA1 across the different brain sections. For vasculature analysis, we adjusted a previously established script that allows the automated calculation of (1) the vascular density, (2) the length of blood vessels, and (3) the number of branches^49,50^. For the permeability analysis, three oval-shaped ROIs (75×80 pixels) were placed outside of blood vessels (stained with CD31) in the stroke core, IBZ, and contralateral hemisphere for normalization. We analyzed three brain slices per mouse and calculated the mean fluorescence intensity of Fibrinogen outside the blood vessels.

### Horizontal ladder walk test

The horizontal ladder walk test was used to identify stroke-related gait abnormalities throughout different timepoints (day 0, 3, 7, 14, 21, 28, 35, 42). Animals were placed on a modified MotoRater 303030 series with a horizontal ladder rung made of metal (length: 113 cm, width: 7 cm) and were trained to cross from one end of the ladder to the other. The structure was made of a clear Plexiglas basin (156 cm long, 11.5 cm wide, 11.5 cm high) and was equipped with two perpendicularly arranged mirrors to allow simultaneous image recording from different angles. To prevent habituation to a specific bar distance, bars were irregularly spaced (1-4 cm) and changed every 2 weeks. At least three runs per animal were recorded with a high-definition video camera (GoPro Hero 12) at a resolution of 4000 x 3000 and a rate of 60 frames per second. Videos were processed in three stages: first, cropped using LosslessCut to isolate frames where the mouse crossed the ladder walk; second, inverted using ShotCut to ensure body-part positions remained consistent during the return journey; and finally, cut with FFmpeg to standardize image coordinates across all filming days. Video recordings were processed by DeepLabCut^TM^ (DLC, v. 3.0.0rc9), an open-source software that enables the computation of 2D and 3D marker-less pose estimates to track limb movements of animals. Initially, 20 frames were extracted from 10 videos and were manually labeled to train the network. Then, we analyzed 30 new videos and extracted outlier frames to refine it. Once the network was adequately trained, recordings of behavioral experiments were fed into DLC for automated tracking. Video pixel coordinates for the labels produced by DLC were imported into R Studio (v. 4.04) and processed with custom scripts that can be assessed here: https://github.com/rustlab1/DLC-Gait-Analysis.

### Statistical analysis

Statistical analysis was performed using R software (v. 4.3.1). For unpaired data, we first evaluated the normality of the data with the Shapiro–Wilk test and the homogeneity of variances with Levene’s test. Based on these diagnostics, for *in vitro* experiments, we used: (1) a one-way ANOVA followed by Tukey’s post-hoc test if the data were normally distributed and variances were homogeneous, or (2) the Kruskal–Wallis test followed by Dunn’s post-hoc test (with Bonferroni correction) if the data were not normally distributed. For *in vivo* experiments, we used: (1) Student’s t-test if the data were normally distributed (p > 0.05) and variances were equal (p > 0.05), (2) Welch’s test if data were normally distributed but variances were unequal (p ≤ 0.05), or (3) Wilcoxon rank-sum test if data was not normally distributed (p ≤ 0.05). For paired data, a linear mixed-effects model (LMM) was fitted with Treatment, Time, and their interaction as fixed effects, and Animal as a random intercept to account for repeated measures. Significance was assessed via Type III ANOVA with Satterthwaite’s degrees of freedom. Post-hoc pairwise comparisons using estimated marginal means (Tukey-adjusted) were conducted only if the Treatment × Time interaction was significant. Data is expressed as mean ± SD (boxplot) and mean ± SEM (ribbon); statistical significance was defined as *p < 0.05, **p < 0.01, ***p < 0.001, and **** p < 0.0001.

## Data availability

All data supporting the findings of this study are available within the article and its supplementary information. Source data for all figures are provided with this paper. Additional data and materials are available from the corresponding author upon request.

## Supporting information

Suupl Tables

## Acknowledgements

This work is supported by the funding from the Mäxi Foundation, Swiss 3R Competence Center (OC-2020-002), Swiss National Science Foundation (SNSF) (CRSK-3_195902) and (PZ00P3_216225) and the Neuroscience Center Zurich.

## Author contributions

B.A.B., R.R., and C.T. designed the study. B.A.B., C.B., C.H., and K.J.Z. planned, conducted, and analyzed *in vitro* experiments. B.A.B., R.Z.W., N.H.R., and C.H. conducted *in vivo* experiments, and B.A.B. and R.R. analyzed them. B.A.B., R.Z.W., and R.R. generated figures. M.G. provided the iPS cell line. R.R. and C.T. supervised the study. B.A.B., N.H.R., R.R., and C.T. wrote the manuscript with input from all authors. All authors read and approved the final manuscript.

