## Supplementary material for "Neural graft elimination via dual safety switch compromises therapeutic recovery after stroke": Suupl Tables

**Supplemental Table 1: Primary Antibody List**

| Antigen | Target | Host | Dilution | Company |
| --- | --- | --- | --- | --- |
| BrdU | Cells dividing at day 25 or 30 (BrdU injection timepoints) | Mouse (IgG2a) | 1:200 | Proteintech |
| HuNu | Human nuclei | Mouse  (IgG1) | 1:200 | EMD Millipore |
| Ki-67 | Marker for cell cycling and proliferation; tumor marker | rabbit | 1:200 | Invitrogen |
| GFAP | Astrocytes | mouse | 1:500 | R&D Systems |
| Iba1 | Microglia | goat | 1:200 | R&D Systems |
| CD 31 | Vascular endothelial cells | rat | 1:50 | BD Biosciences |
| fibrinogen | Blood vessel leakage | rabbit | 1:100 | abcam |

**Supplemental Table 2: Secondary Antibody List**

| Reactivity | Host | Conjugate | Dilution | Company |
| --- | --- | --- | --- | --- |
| Anti-mouse IgG2a | FlexAble | CoraLite 555 | 1:100 | Proteintech |
| Anti-mouse IgG1 | FlexAble | CoraLite 488 | 1:100 | Proteintech |
| Anti-rabbit | Donkey | DyLight405 | 1:200 | *Jackson* |
| Anti-mouse | Donkey | Cy3 | 1:500 | *Jackson* |
| Anti-goat | Donkey | AlexaFluor 647 | 1:500 | *Jackson* |
| Anti-rat | Donkey | Cy3 | 1:500 | *Jackson* |
